# N-of-one differential gene expression without control samples using a deep generative model

**DOI:** 10.1101/2023.01.27.525843

**Authors:** Iñigo Prada-Luengo, Viktoria Schuster, Yuhu Liang, Thilde Terkelsen, Valentina Sora, Anders Krogh

**Affiliations:** Department of Computer Science, University of Copenhagen, Denmark; Center for Health Data Science, University of Copenhagen, Denmark

**Author notes:** equal contributions (Iñigo Prada-Luengo, Viktoria Schuster, Yuhu Liang).

## Abstract

Differential gene expression analysis of bulk RNA sequencing data plays a major role in the diagnosis, prognosis, and understanding of disease. Such analyses are often challenging due to a lack of good controls and the heterogeneous nature of the samples. Here, we present a deep generative model that can replace control samples. The model is trained on RNA-seq data from healthy tissues and learns a low-dimensional representation that clusters tissues very well without supervision. When applied to cancer samples, the model accurately identifies representations close to the tissue of origin. We interpret these inferred representations as the closest normal to the disease samples and use the resulting count distributions to perform differential expression analysis of *single* cancer samples *without* control samples. In a detailed analysis of breast cancer, we demonstrate how our approach finds subtype-specific cancer driver and marker genes with high specificity and greatly outperforms the state-of-the-art method in detecting differentially expressed genes, DESeq2. We further show that the significant genes found using the model are highly enriched within cancer-specific driver genes across different cancer types. Our results show that the *in silico* closest normal provides a more favorable comparison than control samples.

## Introduction

Cellular function varies with cell type and environment and differences in cell function can largely be characterized by gene expression profile (GTEx Consortium 2020; Lonsdale et al. 2013). The analysis of gene expression data has become a standard for studying differences in cells and tissues, partly driven by advances in next-generation RNA sequencing (RNA-Seq) technologies. By identifying differentially expressed genes (DEGs), we can pinpoint genes involved in the onset or progression of disease, which could present biomarkers or potential drug targets in personalized treatment (Burska et al. 2014; Kamel and Al-Amodi 2017).

Despite the great potential of differential expression analysis, the methods employed for this type of analysis yield poorly reproducible results and return thousands of significant DEGs (Cui et al. 2021), making a clinical interpretation challenging. This drawback is largely due to the lack of good controls, a common problem in the study of diseases. In cancer studies, controls are most often tissue samples from healthy individuals (potentially matched on available clinical parameters) or normal adjacent tissues (NATs) from the cancer patients. The benefit of the latter is a reduction of person-specific biological variance. However, NATs from cancer patients have been shown to display some of the same characteristics as the tumor (Aran et al. 2017), implying that these samples are not truly normal. Inversely, normal samples from other individuals will display genetic heterogeneity and are thus unsuitable for direct comparison with classical methods, especially at low sample numbers (D. Li et al. 2022; Vihinen 2022). Lastly, a general problem concerning bulk sequencing data is that samples differ in cell type composition. This problem may in part be alleviated by using weighted averages of the closest normal samples (Rapin et al. 2014; Vivian et al. 2020).

The most generally applied method for differential expression analysis relies on count statistics using negative binomial distributions for accounting for over-dispersion (Love, Huber, and Anders 2014). Neural networks and machine learning in general have been increasingly applied to transcriptomics in the last two decades. Applications range from quality control using simple regression and mixture models (McDermaid et al. 2018) over identifying DEGs and biomarkers using random forests (Abbas and El-Manzalawy 2020) or convolutional neural networks (Kakati et al. 2022) to digital pathology (Schmauch et al. 2020) using multi-layer perceptrons. Other approaches have been suggested to learn biologically meaningful representations from gene expression data. (Way and Greene 2018) present a variational autoencoder (Kingma and Welling 2013) trained on cancer transcriptomes with the potential to predict therapeutic responses. Another generative neural network, SOPHIE (Lee et al. 2022) identifies cancer-specific genes from a collection of normal and cancer datasets. So far, available methods seem to be limited to requiring paired or manually curated controls or training on particular datasets, including cancer samples.

In this work, we present a gene expression model of normal tissue that replaces control samples and enables differential expression analysis in cancer using only a single sample. The model is based on the Deep Generative Decoder (DGD) (Schuster and Krogh 2021, 2022), a generative neural network that learns a probabilistic low-dimensional representation of the data. Our model is trained on the Genotype-Tissue Expression (GTEx) dataset (Lonsdale et al. 2013) containing around 20,000 bulk samples from 31 different human tissues and 948 individuals. Briefly, the model learns parameters with two goals. Firstly, the neural network parameters in the decoder are learned to best describe all data in a low-dimensional space (the “representation”). Secondly, the model learns the most probable representation for each sample in the low-dimensional space and returns a negative binomial distribution over count values for every gene. For samples derived from non-normal tissue, such as cancer samples, we can infer the nearest representation in the model and use it to replace controls. This inferred *in-silico* control is informed by the whole training data instead of a limited and biased set of control samples. Therefore, we expect it to be less noisy and present a more precise judgment of differential expression. To test this hypothesis, we apply the model to cancer samples from The Cancer Genome Atlas (TCGA) program (Cancer Genome Atlas Research Network 2008). From the negative binomial over gene counts, we can derive a p value for each gene in a sample and thus identify a set of significant DEGs. We first focus on analyzing breast cancer (BC) and calculating the enrichment of known cancer driver genes and subtype marker genes among the significant genes. This analysis is compared to a standard case-control analysis using DESeq2, which generally yields many more significant genes and much lower enrichment scores. Finally, we show that cancer driver genes are enriched among the significant genes derived from the DGD across cancers in TCGA.

Altogether, our model of normal gene expression drastically improves differential expression analysis by yielding fewer false positives and extending the analysis to single cancer samples (N-of-one) *without controls*. We believe this method can significantly impact the utility of gene expression analysis, target identification, and personalized treatment.

## Results

Our work aims to construct a deep generative model that learns how genes are expressed across human tissues. Our model, the DGD (Schuster and Krogh 2022), learns a low-dimensional *representation* for every sample. The representation space (or “latent space”) has a dimensionality of 50, and representations are distributed according to a mixture of Gaussians with 45 components. A fully connected feed-forward decoder neural network with two hidden layers maps the latent space to sample space, resulting in a negative binomial distribution for each gene (Fig. 1). We arrived at this setup (decoder architecture, representation dimensionality, and number of Gaussian components) after hyperparameter optimization (see methods). We infer the parameters of the representations, Gaussian mixture model, and decoder by training (Supplementary Fig. S1) our model on a stratified random subset containing 90% of the GTEx data (17072 samples) while leaving the remaining 10% (1903 samples) as a test set (Supplementary Table S1).

**Figure 1:**
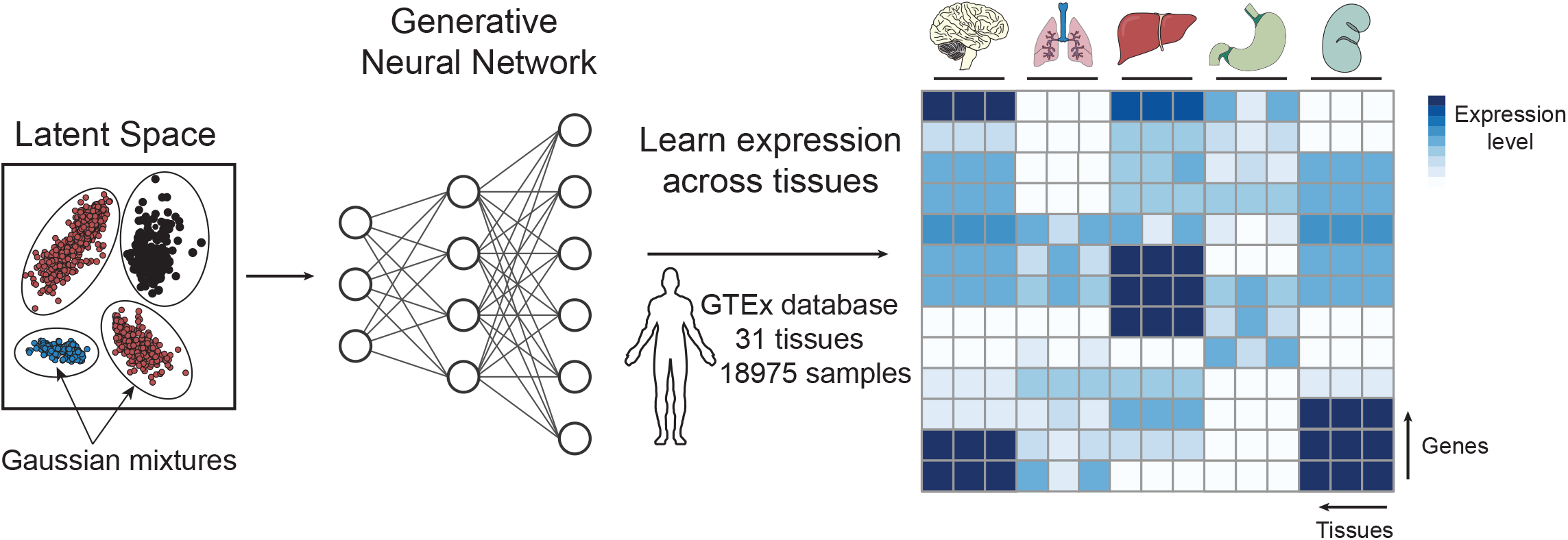
The bulk deep generative decoder. The latent space is parameterized with a Gaussian Mixture Model (left). A generative neural network trained on the GTEx database maps the latent space to the data space. Altogether, the model learns the gene expression distribution across bulk tissues, as illustrated in the heatmap (right).

### A generative model of gene expression for bulk samples

After training, we first evaluated whether the latent space of the DGD model distinguishes different tissues. A two-dimensional principal component analysis (PCA) of the latent space (Fig. 2A) shows that the DGD could find well-separated representations. The learned Gaussian mixture model (GMM) should yield a clustering of tissues based on the mixture components. To test this, we assigned each sample to the GMM component with the highest probability and evaluated how samples are distributed across components. The matrix of associations (Fig. 2B) shows a clear clustering of tissues in the GMM. The clustering of test data (Supplementary Fig. S2) is even clearer, but there are very few samples for some components. A single tissue dominates almost all mixture components. Exceptions to this are two components. These two are split between Uterus and Ovary and Small Intestine and Colon, respectively. Most tissues (20 out of 31) are mainly represented by only one component. Tissues modeled by multiple GMM components are typically split between two or three components. Roughly half of those tissues are represented by multiple subtypes in the data. Interestingly, the brain samples present a clear outlier in the matrix of associations, with this tissue distributed over eight GMM components.

**Figure 2:**
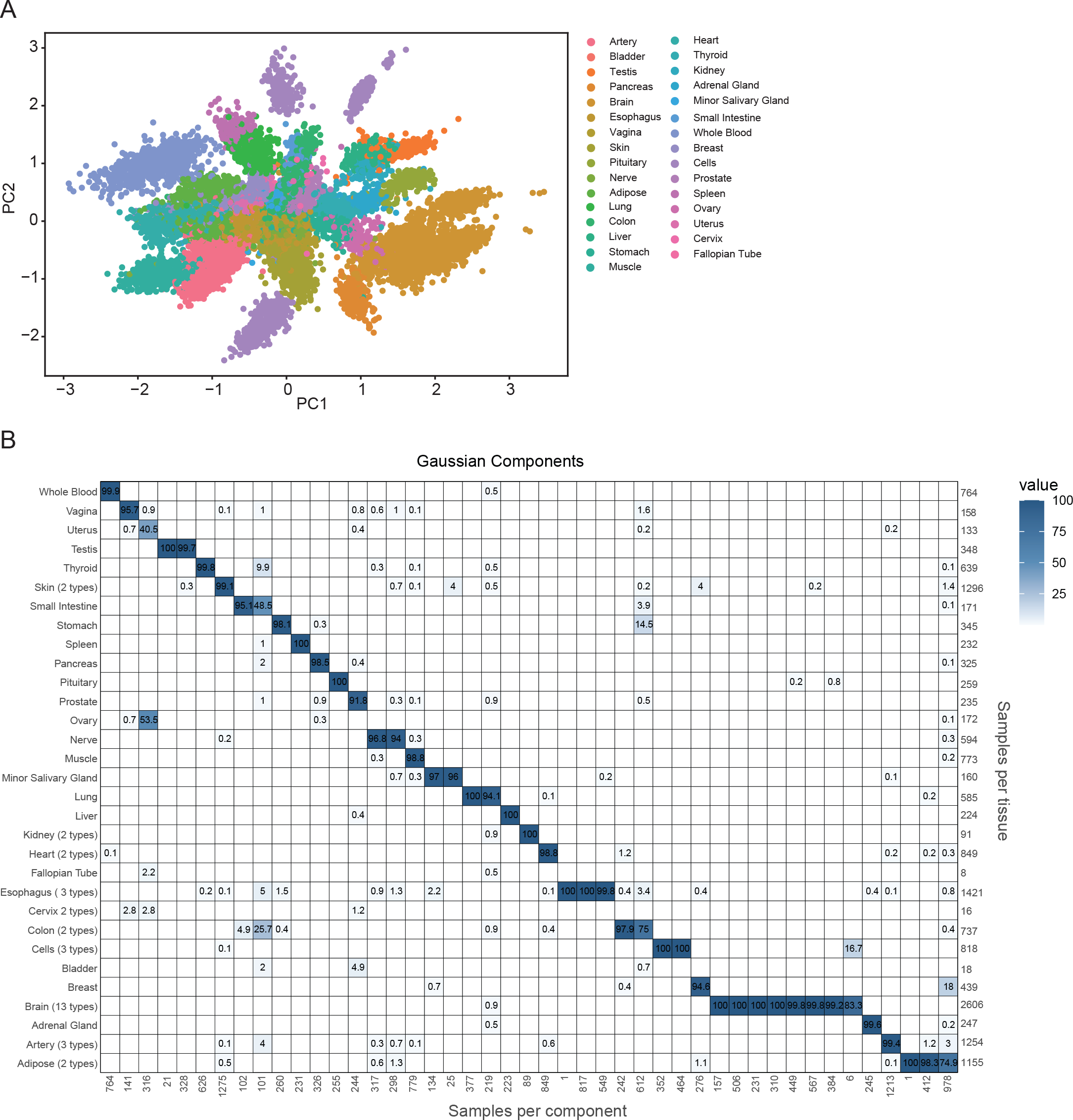
Representations for normal tissue. **A**. PCA plot showing the clustering of 31 tissues in the latent space reduced from 50 dimensions to 2. Each dot corresponds to a training sample. **B**. Matrix of associations showing the percentage of the samples assigned to a Gaussian component (x-axis) represented by each tissue (y-axis). The tissue types and component numbers are sorted for an optimal diagonal view. The numbers below the x-axis give the number of samples assigned to each component. The number of samples in each tissue is shown on the right side of the matrix.

### Finding closest-normal comparison sets for cancer samples

Next, we evaluated whether our model of healthy gene expression could find meaningful representations for cancer samples. To do so, we used our model (trained on GTEx) to find representations for tumor samples (Supplementary Table S2) from The Cancer Genome Atlas dataset (TCGA). These representations were found by maximizing the probability of a representation for a cancer sample while leaving the decoder neural network and GMM parameters fixed (Fig. 3A). We interpreted this representation as the closest-normal sample to the tumor. To begin with, we evaluated the ability of our model to detect out-of-distribution examples (i.e., anomalous expression profiles) by calculating the probability of each sample given our model (Fig. 3B). As observed in Figure 3B, TCGA-cancer samples generally gave a much lower probability than GTEx samples, while TCGA-normal samples achieved probabilities between cancer and GTEx test samples. Afterwards, we assessed whether our model matches tumor samples to the same GMM components as their healthy counterparts (Fig. 3B). We found that our model correctly assigns most tumors to their healthy normal. For 11 out of 14 tissues, more than 80% of the samples are assigned to their corresponding normal tissues. The three tissues with the lowest classification accuracies are Bladder, Stomach, and Esophagus.

**Figure 3:**
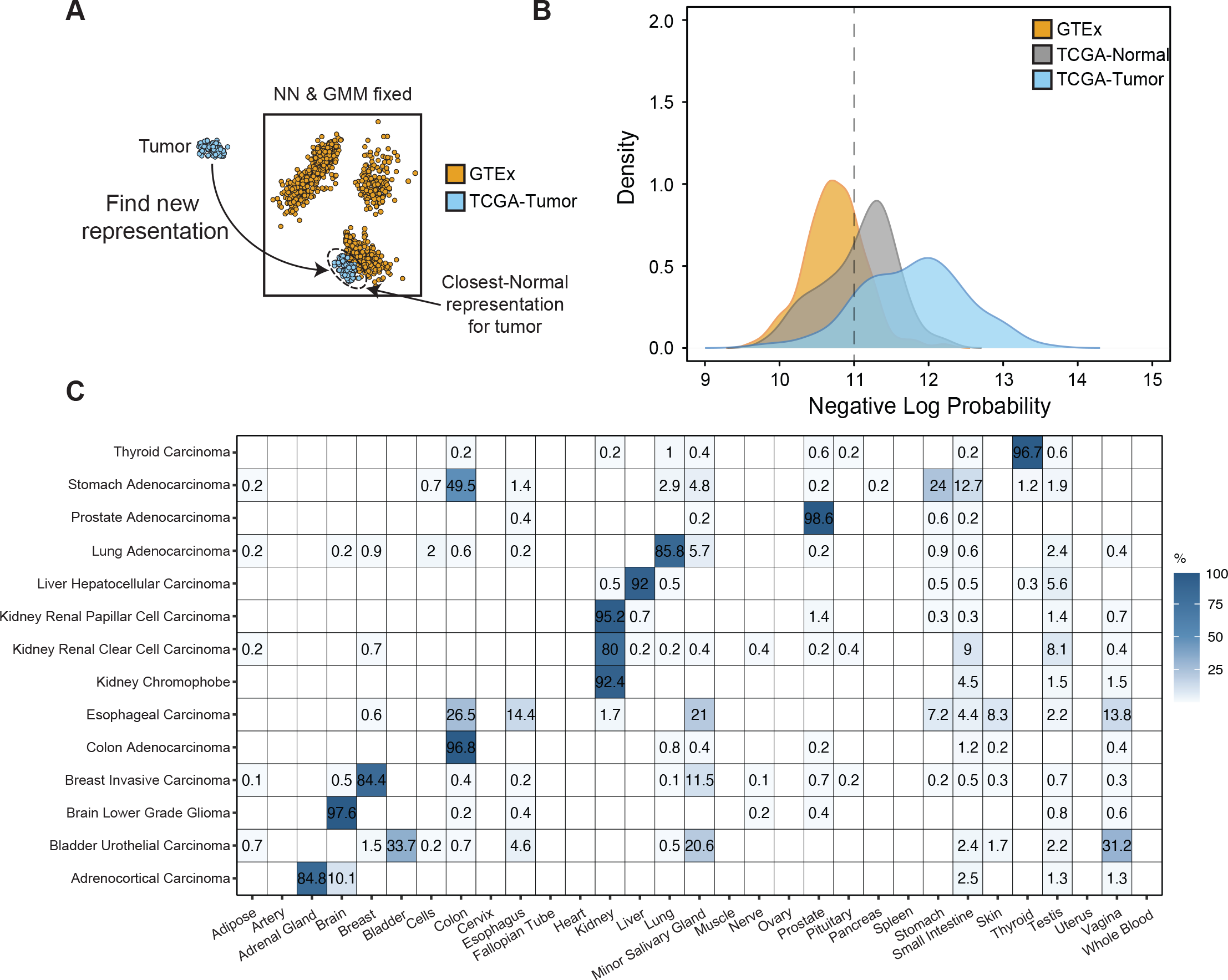
Closest normal representation for cancer samples. **A**. Schematic representation of the procedure to find closest-normal representations for TCGA tumors. We find the representation by maximizing the probability of the representation while leaving the GMM and neural network fixed. **B**. Density plot showing the negative log probability distributions for GTEx test, TCGA-normal and TCGA tumor samples. **C**. Matrix of associations showing the percentage of TCGA tumor samples assigned to GMM components representative of tissues. GMM components are named by their majority normal tissue type.

### Detecting cancer differentially expressed genes without controls

We extended the DGD to detect differentially expressed genes. We performed a two-tailed negative binomial test for the distribution generated from the closest-normal representation (see Material and Methods). Our analysis focused on breast cancer (BC), as both TCGA and GTEx contain a large number of samples (Fig. 4A). We first evaluated the specificity of our approach in a normal vs. normal “negative control” experiment. Afterwards, we tested one cancer sample against the model (N-of-one) to resemble common clinical settings. We compared our approach in both “negative” and “positive control” to a standard analysis of one sample against GTEx normals using DEseq2 (Love, Huber, and Anders 2014), a widely used statistical method to detect differentially expressed genes.

**Figure 4:**
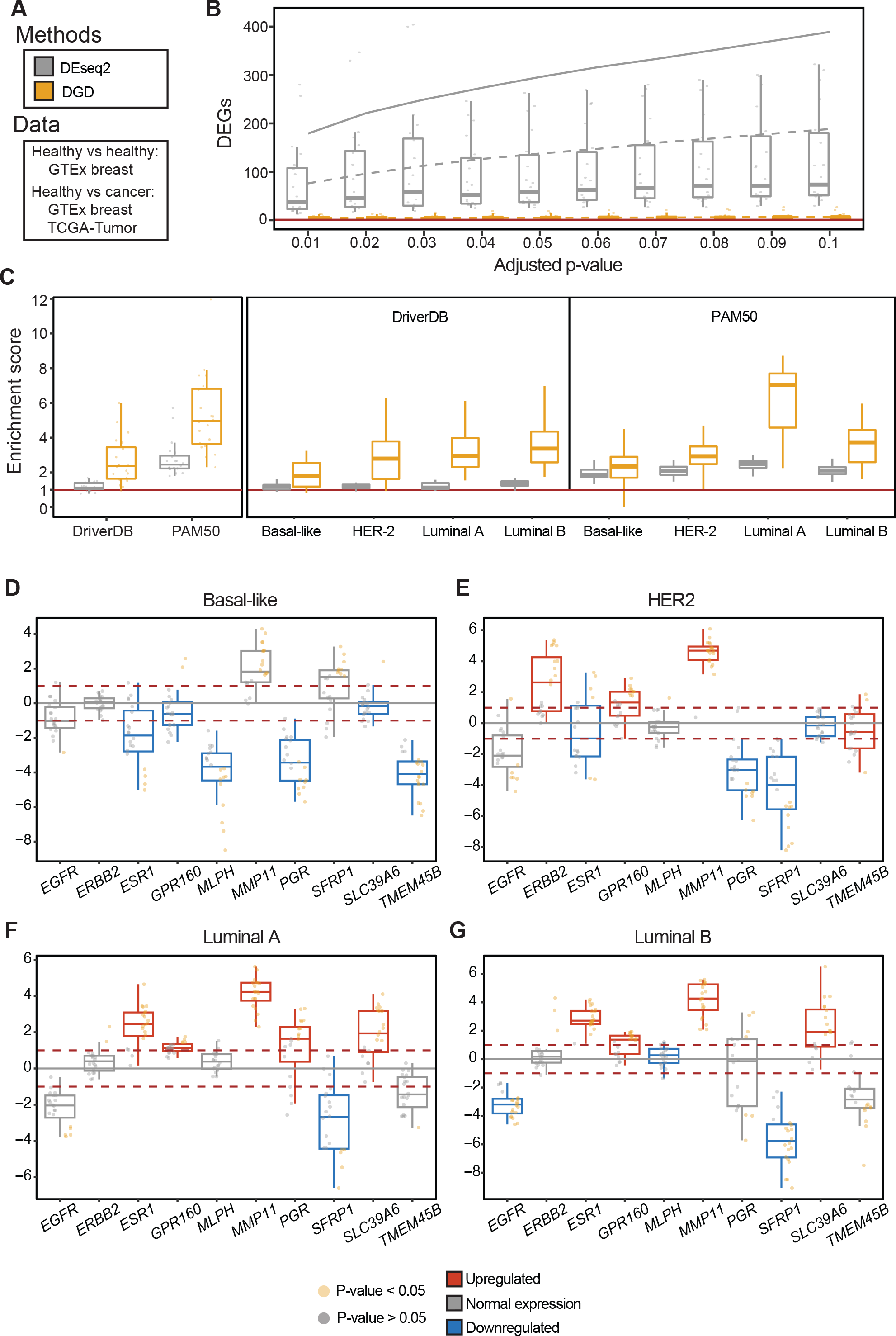
Differential expression analysis of Breast cancer and its subtypes. **A**. Schematic overview of the samples used in our experimental set-up. **B.** “Negative control” experiment comparing healthy test samples against the DGD (yellow) and controls for DESeq2 (gray). 20 random test samples were chosen (shown as dots) and summarized by the boxplots for different cut-offs on adjusted p value (x-axis). As controls for DEseq2, we used both the whole training set of breast tissue (dashed gray line) and a random subset of 5 controls (solid gray line). **C**. Enrichment score across breast cancer subtypes for driver genes and PAM50 genes derived from DGD and DESeq2. 20 random cancer samples were selected and compared to DGD (yellow) and GTEx breast samples as controls for DEseq2 (gray, see Methods for sample selection). The enrichment of cancer driver and PAM50 among the significant genes (adjusted p value < 0.01) is shown for each breast cancer subtype. **D-G** Breast cancer-specific differential expression analysis on a subset of ten marker genes, using 20 randomly selected samples. The box plots are colored based on whether the gene is known to be differentially expressed (red: upregulated; gray: normal expression; blue: downregulated) in a cancer subtype. The dots are colored based on the adjusted p value obtained by DGD in each replicate.

**Figure 5:**
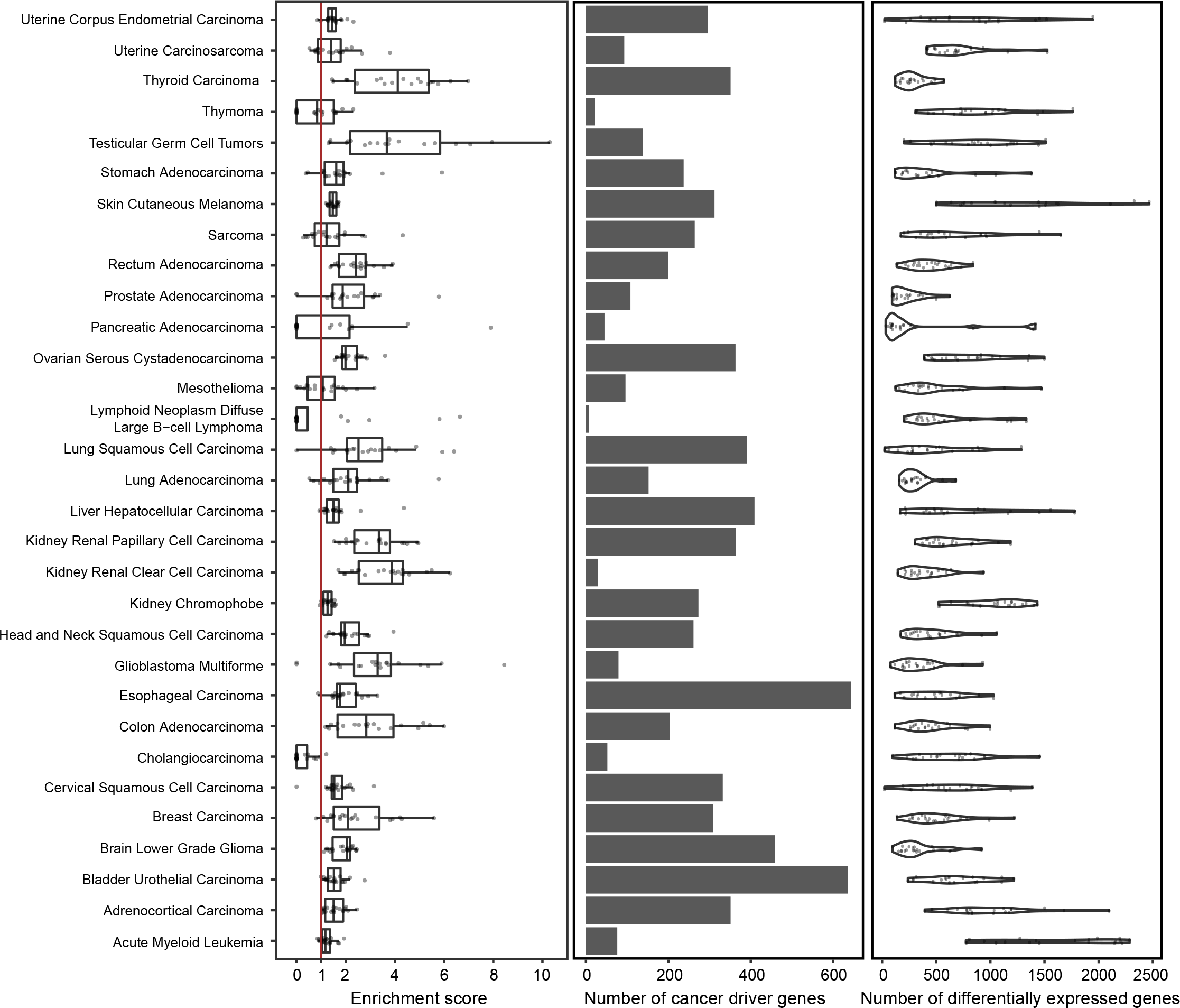
Differential expression and enrichment analysis for cancer driver genes across TCGA. A single sample was randomly selected from each TCGA-tumor type to calculate the differentially expressed genes with our model. This process was repeated 20 times. The left panel shows boxplots of the enrichment scores for every TCGA-tumor. Dots indicate the experiment replicates. The middle plot shows the number of cancer driver genes present in CancerDBv3 for each cancer. The right panel shows a violin plot of the number of differentially expressed genes in every replicate of our experiments.

We assessed the model’s specificity in a normal vs. normal analysis using healthy breast tissue from GTEx. It is assumed that there should be no or very few DEGs when comparing normal samples. Therefore, the number of DEGs here functions as the number of false positives and is thus related to the specificity of the model. We randomly selected 20 samples from the breast test set (out of 42 GTEx samples) and compared each one against the 440 control samples from the training set using DEseq2. We used the same 20 random samples and compared each one to the DGD model. The number of significant genes found by the two methods are shown in Fig. 4B for varying adjusted p values. We performed the same analysis using five randomly selected samples from the training set as the comparison set (full line in Fig. 4B). Ideally, this analysis should return no significant genes. However, DEseq2 found many DEGs, calling on average 76.15 genes (adjusted p value < 0.01 & absolute log2 fold change > 1) using the whole GTEx and 179.05 when using 5 random samples with 20 repetitions as controls. The DGD, on the contrary, found almost no false positives with an average of 4.25 called genes (using the same filtering criteria as DEseq2).

To compare the sensitivities of DGD and DEseq2 in a “positive control” experiment, we analyzed their ability to correctly identify marker genes known to be important in BC. Two sets of BC-related genes were curated for this purpose: (I) driver genes from the DriverDBv3 database (Liu et al. 2020) and (II) the PAM50 (Parker et al. 2009) set of subtype-specific BC genes (Materials and Methods). As a metric, we calculated a gene enrichment score, the fraction of the number of BC-related genes among the significant ones divided by that expected by random chance.

We evaluated enrichment scores across four PAM50 breast cancer subtypes: basal-like, HER2, luminal A, and luminal B. For this analysis, we applied clinical filters to ensure greater homogeneity of samples (Supplementary Table S3). We performed experiments similar to those described above, randomly selecting one sample (20 repetitions) and comparing it to the 143 GTEx breast samples (40-70 year-old females) and the model for DEseq2 and DGD, respectively. The DGD obtained higher enrichment scores than DEseq2 for all subtypes regarding driver genes (DriverDBv3) and PAM50 genes (Fig. 4C). The DGD obtained an average enrichment score of 3.46 and attained a particularly high score for the luminal A subtype in the PAM50 marker set. The DEseq2 average score was 1.71, and it was lower than the model for subtypes (note that a score of 1 means no enrichment).

Next, we selected a subset of PAM50 genes to evaluate whether the DGD captured the expression differences between subtypes. We focused on genes that achieved a significant p value in at least 10 replicates of the PAM50 experiment performed above(Supplementary Table S4), leading to ten genes across the four subtypes. For each gene, we evaluated whether the DGD detected differential expression (i.e. a significant p value) and the expression trend (i.e., upregulation or downregulation) as defined in the PAM50 set (Coleman and Anders 2017; Vaca-Paniagua et al. 2015; Hu et al. 2013; Gendoo et al. 2016). The DGD correctly identified 32 of the 40 expected gene expression patterns (Fig 4. D-G), obtaining similar performances across the subtypes. The best overall performance was achieved for the luminal A and luminal B subtypes (eight out of ten patterns determined correctly). The lowest performance was for basal-like and HER2 subtypes (six out of ten genes correctly determined). Summarizing across genes, *ERBB2*, *GPR160*, and *PGR* they have the expected trend in all subtypes. *ESR1, SFRP1, MLPH*, and *MMP11* have the expected trend in three out of four subtypes. The DGD only under-performed on *EGFR, SLC39A6*, and *TMEM45B*, which were wrongly identified in two out of four, two out of four and three out of four subtypes, respectively.

As a final analysis, we extended our model to all the 31 TCGA cancer types to better estimate the DGD’s general performance. As described above, we performed N-of-one experiments using 20 repetitions, and we evaluated the enrichment score of the DriverDBv3 gene set for each cancer type (Supplementary Table S5). The DGD found more cancer marker genes than expected by chance for all cancer types (mean enrichment score 2.05, mean range 0.24-4.70). Specifically, the scores were higher for Kidney Renal Clear and Papillary Cell Carcinomas (mean enrichment of 3.65 and 3.18, respectively),Thyroid Carcinoma (mean enrichment 4.00), and Testicular Germ Cell Tumors (mean enrichment 4.70). Cancers with median enrichment below 1 tend to have a small number of cancer driver genes in the database (Thymoma, Pancreatic Adenocarcinoma, Lymphoid Neoplasm Diffuse Large B−cell Lymphoma and Cholangiocarcinoma).

## Discussion

A lot of attention is currently directed towards single-cell RNA sequencing due to its potential higher resolution and recent advances in its scalability. However, bulk sequencing is still a work-horse for clinical use (X. Li and Wang 2021). Differential expression analysis between disease and normal has relied on solid statistical methods but is challenged by the difficulty in obtaining suitable control samples and large enough sample sizes. Here, we introduce a method requiring no biological replicates or matched controls. The model we present learns the gene expression of normal tissue samples and generates an *in silico* normal sample closest to the disease sample at hand.

To assess whether the model learns meaningful representations, we analyzed how well GMM components model tissues as the most prevalent biological feature in the data. The model clusters representations well in a tissue-dependent manner and typically assigns only one to a few Gaussian mixture components to each tissue type. Most tissues not being modeled by a single component are represented by multiple subtypes in the data or present complex biological tissues, such as the brain. Since the number of components in the GMM is much higher than the number of tissues, it is understandable that multiple components model more complex structures. It also makes sense that under-represented tissues from larger systems are clustered together. An example is the component gathering samples from Uterus, Ovary, Fallopian Tube, and Cervix. Given that the representations are found in a completely unsupervised fashion, we find the clustering to be remarkably interpretable.

Besides providing well-clustered representations, the model of normal gene expression can replace control samples in the differential gene analysis of disease. This is achieved by finding the closest normal representation in the model. This representation is used to compute a probability distribution for the expression counts of each gene. We calculated the total probabilities of the GTEx test, TCGA normal, and TCGA cancer samples. We found that the probabilities derived from the model of normal are highest for GTEx test data and decrease strongly for TCGA cancer as expected. Probabilities for TCGA normal samples lie between the two, which may be explained by previous findings suggesting that adjacent normal tissue carries cancer traits (Aran et al. 2017). A similar analysis has previously been performed using a Bayesian approach (Vivian et al. 2020), but the separation seems much clearer (Vivian et al. 2020), although a direct comparison is difficult. When analyzing the quality of the integration of cancer samples into latent space, we observe that most representations of cancer samples are consistent with the tissue of origin. For most cancers, more than 80% of samples end up in the expected tissue component. These results are comparable to (Vivian et al. 2020). Here it is worth noting that we do not necessarily expect a perfect match to the tissue of origin because, in metastatic cancers, the tumor may be more similar to the tissue of origin.

It is difficult to assess the performance of differential gene expression analysis without knowing the ground truth. We have therefore used an enrichment score for known cancer genes among the significant genes to compare our model to a standard analysis using breast cancer as a case. Firstly, we assessed the specificity of our model and DESeq2 by comparing normal vs. normal as a negative control. While DESeq2 yields large numbers of significant genes (1.75% of all genes, given a log2-fold change > 1 and an adjusted p value < 0.05), here interpreted as false positives, our model reports only 0.03% of the genes to be differentially expressed under the same conditions. Secondly, we calculate the enrichment of relevant known cancer genes among the significant genes derived from differential expression analysis of breast cancer samples. In this positive control, we used a set of known breast cancer driver genes and a PAM50 set of genes discriminating between breast cancer subtypes. In the general breast cancer case, we see consistently higher enrichment when using the DGD compared to DESeq2. This is also true, for the most part, for the cancer subtypes. However, there are some discrepancies with respect to the expected driver genes. For instance, according to our model, *TMEM45B* should be upregulated in the *HER2* subtype but is downregulated on average. Yet, we do not expect a perfect concordance between the PAM50 panel and tumor samples, as many patient-specific factors can affect gene expression. Altogether, our evaluation of breast cancer shows that the DGD returns much fewer false positives and a higher proportion of truly relevant genes compared to DESeq2.

The model introduced and discussed here presents an important step towards using bulk gene expression analysis for precision medicine. Given that the DGD does not require control samples, the potential impact for differential expression analysis with *single disease samples* is immense. We have tested the method on cancer, but there is no reason that it should not perform as well on other diseases. Because of the impressive performance on even single samples, we believe our model has a high potential for application to rare diseases. Since the differential expression analysis generally results in much fewer false positives than in a standard analysis, it will enable us to find genes that are truly involved in disease. This can in turn, increase the possibility to find druggable targets and to understand diseases on an individual level much better than it has previously been possible with bulk data.

## Methods

### The model

#### Architecture and hyperparameters

The full model consists of a Gaussian mixture model as the parametrized distribution over latent space, and a decoder as presented in (Schuster and Krogh 2022). We performed hyperparameter optimization on the latent dimensionality, number of hidden layers in the decoder, and number of Gaussian components. Representations were explored with dimensionalities between 10 and 100. We further tested various decoder architectures with two to four hidden layers of 200 to 10000 hidden units. For the Gaussian mixture model, we tried different distribution complexities with 20 to 50 components. Each representation of a sample received a 50-dimensional vector initialized with zero. The decoder consists of a 50-dimensional input layer fed with the representations, two hidden layers, and an output layer with units corresponding to each of the genes in the data. The two hidden layers are of size 500 and 8000, respectively, immediately followed by a ReLU activation (Fukushima 1975; “Rectified Linear Units Improve Restricted Boltzmann Machines” n.d.). The output values are transformed into negative binomial distributions over expression counts for each gene. The negative binomial is given by:

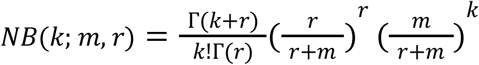

where *k* is the library size, *m* is the mean and *r* the dispersion parameter. The decoder outputs are passed through ReLU activation and scaled with the sample gene expression mean and used as the mean (m) of the distributions. The trainable dispersion parameters (r) of the negative binomials are gene-specific and initialized with 2. The GMM consists of 45 mixture components. The priors are a mollified uniform distribution (the “softball” prior) with a spread of 7 and a sharpness of 10 for the means, a Gaussian with a mean of 1 (corresponds to a standard deviation of 0.1) and a standard deviation of 1 for the negative logarithmic diagonal covariance and a Dirichlet distribution with all alphas equal to five. For more details, please refer to (Schuster and Krogh 2022).

#### Training

The model was trained for 200 epochs with a batch size of 256. The optimizer of choice was Adam (Kingma and Ba 2014) without weight decay and betas 0.5 and 0.9. Because the representations are updated every epoch, the decoder, representation and GMM received distinct optimizer instances with learning rates 10^−4^, 0.01 and 0.01, respectively.

#### Evaluation

Representations for single test samples are optimized as described in (Schuster and Krogh 2022) with all model parameters fixed. For a new datapoint, representations were initialized from the component means. This results in 45 representations per sample. These were optimized with the fixed model for 20 epochs, after which the best representation is selected for each sample and optimized for another 50 epochs.

#### Differential expression

We used the DGD to find differentially expressed genes for tumor samples. An optimal representation is found for each tumor sample as described above. These representations are interpreted as the closest-normal for each tumor – an *in silico* normal. In our setup, we want to test if counts for a given gene are significantly different between the tumor sample and the normal tissue output of the neural network. Let *NB*(*k*; *m_i_*, *r_i_*) be the negative binomial distribution for gene *i* and *x_i_* be the actual count in the tumor sample. We test the hypothesis that the observation from the tumor is from the *in silico* normal. Therefore, the p value for *x_i_* is the sum of all counts with a probability lower than that of *x_i_* :

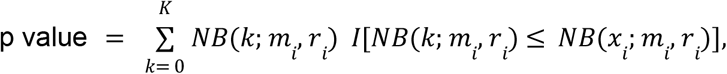

where *K* is the library size and *I* is an indicator function taking a value of 1 when the condition is fulfilled and otherwise 0. The above expression yields an exact p value for the negative binomial distributions. However, it requires summing over all read counts across genes. For the sake of efficiency, we therefore obtain an asymptotic p value by summing over an evenly spaced grid of 10^4^ in the domain of *K*.

### Data

#### Data collection and processing

The raw gene count expression data from the Genotype-Tissue Expression (GTEx) and The Cancer Genome Atlas (TCGA) were downloaded from the Recount3 database (https://rna.recount.bio/), using the built-in R packages. Additionally, the sample metadata files were acquired from the Recount3 platform (Wilks et al. 2021).

At the time of download (Feb 9th, 2022) there were 31 different tissue types in GTEx and one group without tissue information, with a total of 19,214 individual samples (Supplementary Table S1). The 133 samples without tissue information were removed, and duplicates were dropped before further processing. We performed feature selection using the *filterbyExpr* (Chen, Lun, and Smyth 2016) function with default parameters in R (edgeR version 3.34.1) to remove lowly expressed genes. We further reduced the set of genes to protein-coding genes according to (Karolchik et al. 2004). After preprocessing, the GTEx dataset contained 18,975 samples and 16,883 annotated protein-coding genes.

For training and evaluation, the samples were split into a train and a test set, with the test set representing roughly 10% of the data. Sampling was performed in a proportional stratified random manner, meaning that 10% of the samples were randomly chosen and assigned to the test set for each tissue. This procedure resulted in 17072 train and 1903 test samples.

As for cancer data, The Cancer Genome Atlas (TCGA) Program provided 33 different cancer types. We selected the set of genes present in our processed GTEx dataset and separated the TCGA samples into a normal adjacent set (747 samples) and a tumor set (10601 samples) following the metadata file.

#### TCGA tissue selection

For an assessment of the model’s capability to assign independent samples to the correct tissue clusters according to the training data, we selected 14 cancers from TCGA that fulfilled the following two conditions: 1) the tissue corresponding to the cancer is present in GTEx, and 2) the tissue must have at least ten adjacent normal samples from each cancer type (Zeng et al. 2019). We also included the Adrenal and Brain tumor samples, even though they lack adjacent normal samples, to compare our result with the work of Vivian et al. (Vivian et al. 2020). This results in 6111 TCGA tumor samples and 624 TCGA adjacent normal samples (Supplementary Table S2). In applying our differential expression analysis to cancer types other than breast cancer, we include TCGA cancer samples from the 31 cancer types overlapping with those found in DriverDBv3 (Liu et al. 2020).

#### TCGA breast cancer subset

To obtain a homogenous breast cancer (BC) dataset for testing, we curated the TCGA BC samples to only include primary tumors from women between 40 and 70 years of age. We excluded samples with low tumor cell percentage (defined as < 50%) or a high level of necrosis (defined as > 5 %), in addition to samples from patients with known metastasis, stage IV or stage X tumors, or prior cancer diagnosis. For a full list of selection criteria and columns from metadata used for curation, see Supplementary Table S4. The number of available BC samples for analysis was reduced from 1256 to 381.

#### Cancer Driver Genes and PAM50 genes

We downloaded a list of cancer driver genes for each cancer type from DriverDBv3 (Liu et al. 2020) and we matched the symbol names in the database to the Ensembl IDs from recount3. The PAM50 gene set used for BC subtype classification (Basal, Luminal A, Luminal B and Her2-enriched) (Parker et al. 2009) was downloaded through the R-package *genefu (Gendoo et al. 2016)*. As our model testing and comparison with DEseq2 pertained to the expression profile of BC subtype tissue vs. normal tissue (i.e., not between subtypes), we filtered the PAM50 dataset to only include the genes which were specific to a single subtype or which could distinguish BC subtype(s) from normal tissue. We noted the expected directionality of each of the PAM50 genes (up- or down-regulated) for a contrast. The selection of genes was based partly on literature (Coleman and Anders 2017; Vaca-Paniagua et al. 2015; Hu et al. 2013) and partly on the robust normalized PAM50 scores (Gendoo et al. 2016). The PAM50 gene set was reduced to 34 genes.

### Analysis

#### Evaluation of tissue specificity

Clustering performance according to tissue type was evaluated based on the GMM probability densities for each sample’s representation. For this purpose, samples are assigned the GMM component that achieves the highest probability density for their inferred representations. We calculated the percentage of each tissue per component as the number of samples of a given tissue clustered in a given component divided by the total number of samples assigned to this component.

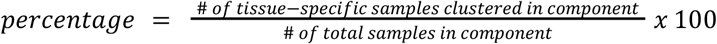

This is done independently for the train set (Fig. 2B) and test set (Fig S2).

#### Matching TCGA to normal tissues

We use the TCGA data described above to evaluate the mapping of unseen and independent data onto the latent space. New representations for all 6111 TCGA tumor samples were found using the DGD trained on GTEx data. The resulting GMM probability densities for the TCGA representations are used to evaluate how well new samples match the correct tissues of the training representation. We define the “correct” tissue as the tissue that best represents a given GMM component. Our evaluation metric is the percentage of TCGA samples of a given tissue matched to the corresponding GMM component with respect to the total number of TCGA samples for this tissue.

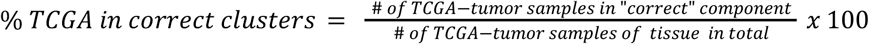

Bladder samples were evaluated differently due to the lack of a bladder-specific component in the normal model. Instead, we evaluated a correct match as TCGA and GTEx bladder samples were assigned to the same component(s).

#### Comparing GTEx and TCGA gene expression predictions

The total probability of a sample is the product of the probabilities across all 16,883 genes given by the model (also used in the model likelihood). We calculate the negative log-probability mass of each sample for three datasets: GTEx test, TCGA adjacent normal and TCGA tumor. For this comparison, we use subsets containing ten tissues: Adrenal, Brain, Breast, Colon, Kidney, Liver, Lung, Prostate, Stomach, and Thyroid. We apply this analysis for a pan-tissue comparison based on the eight tissues only, because Adrenal and Brain are missing in the TCGA adjacent normal subset. For a fair comparison, we ensure equal samples for a given tissue across the three datasets. If all datasets have more than 20 samples for a given tissue, we randomly select 20 samples from each subset for that tissue.

#### Differential expression analysis in TCGA Breast Cancer

Differential expression analysis (DEA) performed by our model is compared to DESeq2 by theenrichment of cancer driver genes among DEGs.

For a general comparison, we perform 20 single-sample experiments using 1 random breast cancer sample (case) from the population of 40-50 year-old caucasian females. This procedure leaves us with 166 samples. Genes that result in absolute log2-fold changes greater than one and adjusted p values below 0.01 are accepted as differentially expressed. The enrichment score is then given as the normalized number of DEGs belonging to the group of breast cancer driver genes or PAM50 genes, respectively.

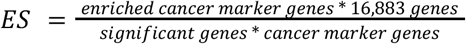

We perform a comparable DEA with DESeq2 using all 44 GTEx samples (control) under the same conditions (40-50 year-old females).

We also perform single-sample analyses of the four available breast cancer subtypes: Basal-like (84 samples), HER2 (37 samples), Luminal A (176 samples), and Luminal B (84 samples), both for our model and DEseq2. We randomly choose one sample from each of the four subtypes as a case sample and used all GTEx breast tissue samples (40-70 year-old females, 143 samples in total) as controls in the DEseq2 method. The experiment is repeated 20 times for each subtype.

#### False positive analysis

In order to assess the false positive rates of the methods, we selected a random GTEx breast sample from the test set (42 samples) as a false case sample. We performed DEA with our model and DESeq2 to yield false positive DEGs for adjusted p values ranging from 0.01 to 0.1 (absolute log2-fold change greater than 1). We performed this 20 times and reported the resulting DEGs as false positives. As controls for DESeq2, we used all breast samples from the GTEx training set (440 samples) as controls. We also perform the analysis for DEseq2 by choosing five controls, which are randomly selected from the breast samples mentioned above.

For the enrichment analysis of Cancer Driver Genes for multiple cancer types, 31 different cancer types are used: Adrenocortical Carcinoma, Bladder Urothelial Carcinoma, Brain lower Grade Glioma, Breast Carcinoma, Colon Adenocarcinoma, Kidney cancer (Kidney Chromophobe, Kidney Renal Clear Cell Carcinoma, Kidney Renal Papillary Cell Carcinoma), Liver Hepatocellular Carcinoma, Lung Adenocarcinoma, Prostate Adenocarcinoma, Stomach Adenocarcinoma, and Thyroid Carcinoma. We perform 20 single-sample experiments for each of the cancer types. For each cancer type, its respective cancer driver gene list was used in the enrichment score calculation.

## Code Availability

The code and our model are available on GitHub: https://github.com/Center-for-Health-Data-Science/BulkDGD.

## Acknowledgements

We would like to acknowledge discussions with Jonas Sibbesen and the Center for Health Data Science and the Center for Basic Machine Learning Research in Life Science in general.

## Supplementary Figures

**Supplementary Figure S1:**
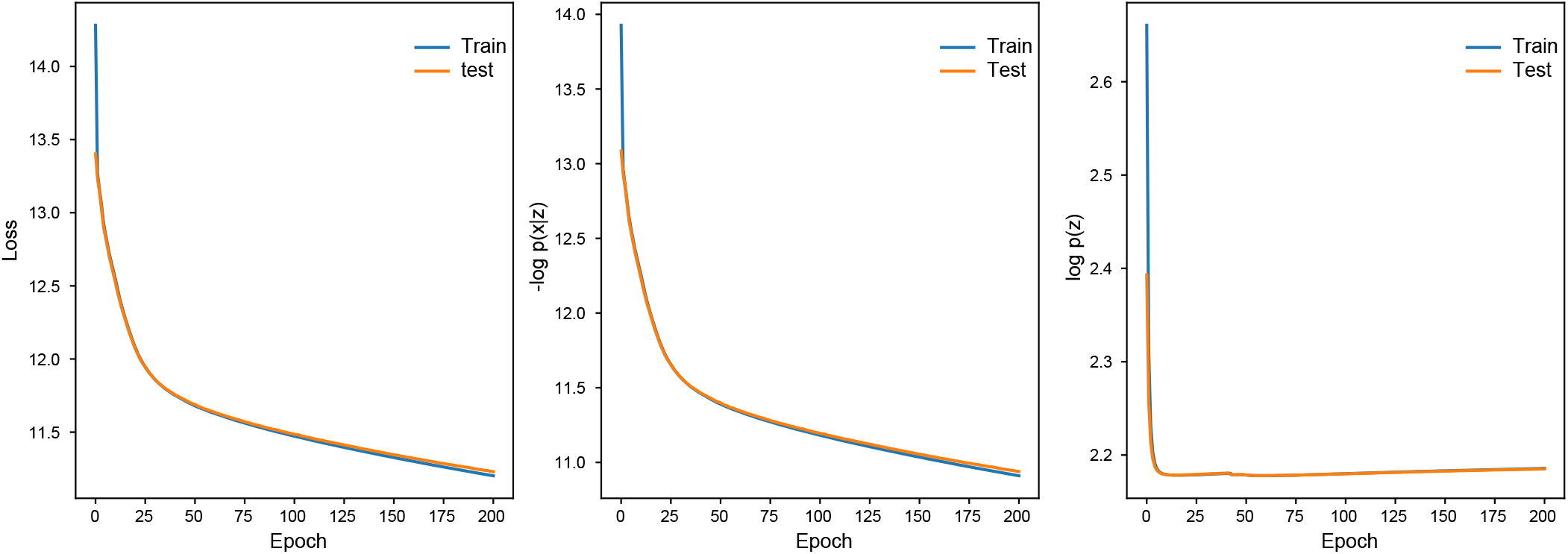
Loss curves. The figure shows loss curves for training and test sets in blue and orange, respectively. The left panel shows the total loss curve for the DGD model. The two panels on the right separate the loss into two terms. The middle panel shows the reconstruction loss of the DGD, and the right panel shows the Gaussian mixture model loss.

**Supplementary Figure S2:**
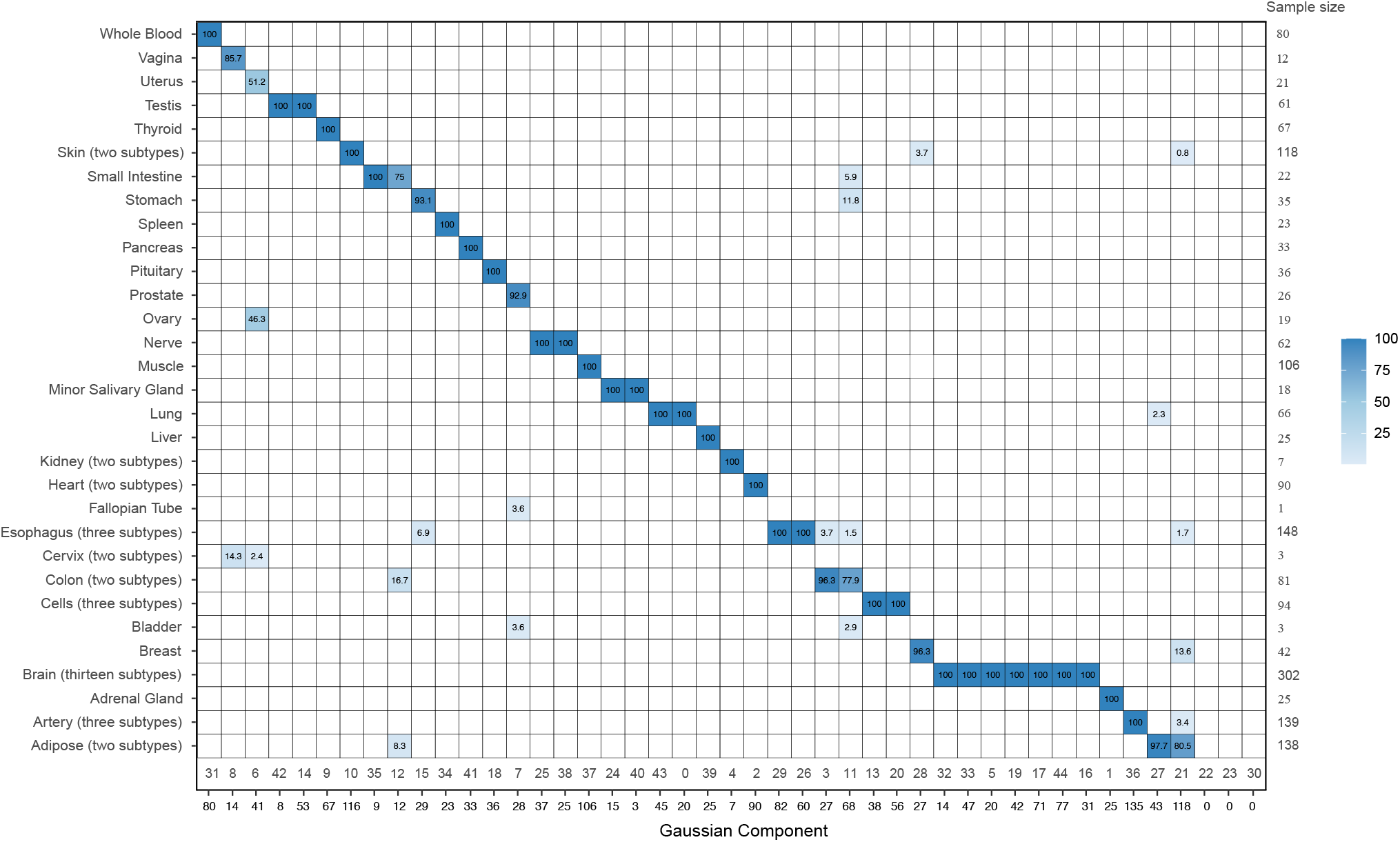
Matrix of associations for the GTEx test set. The matrix shows the percentage of samples assigned to the Gaussian components (x-axis) represented by each tissue (y-axis). The tissue types and components have the same order as in Figure 2B. The row inmediately below the matrix shows the indexes of the Gaussian components. The row below shows the number of samples in each component. The column on the right shows the number of samples in each tissue.

**Supplementary Table S1. Overview of the samples from GTEx**. 31 different tissue sites are represented in the table with their corresponding sample sizes. The samples were downloaded from the Recount3 platform. The last two columns of the table represent the number of samples used in the training and test sets, respectively.

**Supplementary Table S2. Overview of TCGA tumor and TCGA adjacent samples**. The table presents the 31 cancer types (TCGA normal and adjacent) used in our study. The first 14 cancers are the selected ones (see Methods) used to evaluate whether the DGD finds correct comparison samples (Fig. 3C). All the cancers are used in Fig. 5C.

**Supplementary Table S3. Clinical filters for selecting breast cancer subtypes**. The table shows the clinical criteria we used to select samples from the four different subtypes of breast cancer. We downloaded the metadata associated with the samples from the Recount3 platform and applied the filters.

**Supplementary Table S4. 34 PAM50 genes**. The table contains the expected expression trend (downregulated or upregulated) for the selected PAM50 genes. The last four columns represent the number of times the DGD model identified each gene as significant. Criteria for significance were a p-value lower than 0.05 and an absolute log2 fold change greater than 1.

**Supplementary Table S5. Number of cancer driver genes**. This table shows the number of driver genes for each cancer type. The driver genes are obtained from DiverDBv3 (Methods).

## References

Abbas, Mostafa, and Yasser El-Manzalawy. 2020. “Machine Learning Based Refined Differential Gene Expression Analysis of Pediatric Sepsis.” BMC Medical Genomics 13 (1): 122.

Aran, Dvir, Roman Camarda, Justin Odegaard, Hyojung Paik, Boris Oskotsky, Gregor Krings, Andrei Goga, Marina Sirota, and Atul J. Butte. 2017. “Comprehensive Analysis of Normal Adjacent to Tumor Transcriptomes.” Nature Communications. https://doi.org/10.1038/s41467-017-01027-z.

Burska, A. N., K. Roget, M. Blits, L. Soto Gomez, F. van de Loo, L. D. Hazelwood, C. L. Verweij, et al. 2014. “Gene Expression Analysis in RA: Towards Personalized Medicine.” The Pharmacogenomics Journal 14 (2): 93–106.

Cancer Genome Atlas Research Network. 2008. “Comprehensive Genomic Characterization Defines Human Glioblastoma Genes and Core Pathways.” Nature 455 (7216): 1061–68.

Chen, Yunshun, Aaron T. L. Lun, and Gordon K. Smyth. 2016. “From Reads to Genes to Pathways: Differential Expression Analysis of RNA-Seq Experiments Using Rsubread and the edgeR Quasi-Likelihood Pipeline.” F1000Research 5 (June): 1438.

Coleman, William B., and Carey K. Anders. 2017. “Discerning Clinical Responses in Breast Cancer Based On Molecular Signatures.” The American Journal of Pathology 187 (10): 2199–2207.

Cui, Weitong, Huaru Xue, Lei Wei, Jinghua Jin, Xuewen Tian, and Qinglu Wang. 2021. “High Heterogeneity Undermines Generalization of Differential Expression Results in RNA-Seq Analysis.” Human Genomics 15 (1): 7.

Fukushima, K. 1975. “Cognitron: A Self-Organizing Multilayered Neural Network.” Biological Cybernetics 20 (3-4): 121–36.

Gendoo, Deena M. A., Natchar Ratanasirigulchai, Markus S. Schröder, Laia Paré, Joel S. Parker, Aleix Prat, and Benjamin Haibe-Kains. 2016. “Genefu: An R/Bioconductor Package for Computation of Gene Expression-Based Signatures in Breast Cancer.” Bioinformatics 32 (7): 1097–99.

GTEx Consortium. 2020. “The GTEx Consortium Atlas of Genetic Regulatory Effects across Human Tissues.” Science 369 (6509): 1318–30.

Hu, Ying, Ling Bai, Thomas Geiger, Natalie Goldberger, Renard C. Walker, Jeffery E. Green, Lalage M. Wakefield, and Kent W. Hunter. 2013. “Genetic Background May Contribute to PAM50 Gene Expression Breast Cancer Subtype Assignments.” PloS One 8 (8): e72287.

Kakati, Tulika, Dhruba K. Bhattacharyya, Jugal K. Kalita, and Trina M. Norden-Krichmar. 2022. “DEGnext: Classification of Differentially Expressed Genes from RNA-Seq Data Using a Convolutional Neural Network with Transfer Learning.” BMC Bioinformatics. https://doi.org/10.1186/s12859-021-04527-4.

Kamel, Hala Fawzy Mohamed, and Hiba Saeed A. Bagader Al-Amodi. 2017. “Exploitation of Gene Expression and Cancer Biomarkers in Paving the Path to Era of Personalized Medicine.” Genomics, Proteomics & Bioinformatics 15 (4): 220–35.

Karolchik, Donna, Angela S. Hinrichs, Terrence S. Furey, Krishna M. Roskin, Charles W. Sugnet, David Haussler, and W. James Kent. 2004. “The UCSC Table Browser Data Retrieval Tool.” Nucleic Acids Research 32 (Database issue): D493–96.

Kingma, Diederik P., and Jimmy Ba. 2014. “Adam: A Method for Stochastic Optimization.” arXiv [cs.LG]. arXiv. http://arxiv.org/abs/1412.6980.

Kingma, Diederik P., and Max Welling. 2013. “Auto-Encoding Variational Bayes.” arXiv [stat.ML]. arXiv. http://arxiv.org/abs/1312.6114v10.

Lee, Alexandra J., Dallas L. Mould, Jake Crawford, Dongbo Hu, Rani K. Powers, Georgia Doing, James C. Costello, Deborah A. Hogan, and Casey S. Greene. 2022. “SOPHIE: Generative Neural Networks Separate Common and Specific Transcriptional Responses.” Genomics, Proteomics & Bioinformatics, October. https://doi.org/10.1016/j.gpb.2022.09.011.

Li, Dongmei, Martin S. Zand, Timothy D. Dye, Maciej L. Goniewicz, Irfan Rahman, and Zidian Xie. 2022. “An Evaluation of RNA-Seq Differential Analysis Methods.” PloS One 17 (9): e0264246.

Liu, Shu-Hsuan, Pei-Chun Shen, Chen-Yang Chen, An-Ni Hsu, Yi-Chun Cho, Yo-Liang Lai, Fang-Hsin Chen, et al. 2020. “DriverDBv3: A Multi-Omics Database for Cancer Driver Gene Research.” Nucleic Acids Research 48 (D1): D863–70.

Li, Xinmin, and Cun-Yu Wang. 2021. “From Bulk, Single-Cell to Spatial RNA Sequencing.” International Journal of Oral Science 13 (1): 36.

Lonsdale, John, Jeffrey Thomas, Mike Salvatore, Rebecca Phillips, Edmund Lo, Saboor Shad, Richard Hasz, et al. 2013. “The Genotype-Tissue Expression (GTEx) Project.” Nature Genetics 45 (6): 580–85.

Love, Michael I., Wolfgang Huber, and Simon Anders. 2014. “Moderated Estimation of Fold Change and Dispersion for RNA-Seq Data with DESeq2.” Genome Biology 15 (12): 550.

McDermaid, Adam, Xin Chen, Yiran Zhang, Cankun Wang, Shaopeng Gu, Juan Xie, and Qin Ma. 2018. “A New Machine Learning-Based Framework for Mapping Uncertainty Analysis in RNA-Seq Read Alignment and Gene Expression Estimation.” Frontiers in Genetics 9 (August): 313.

Parker, Joel S., Michael Mullins, Maggie C. U. Cheang, Samuel Leung, David Voduc, Tammi Vickery, Sherri Davies, et al. 2009. “Supervised Risk Predictor of Breast Cancer Based on Intrinsic Subtypes.” Journal of Clinical Oncology: Official Journal of the American Society of Clinical Oncology 27 (8): 1160–67.

Rapin, Nicolas, Frederik Otzen Bagger, Johan Jendholm, Helena Mora-Jensen, Anders Krogh, Alexander Kohlmann, Christian Thiede, et al. 2014. “Comparing Cancer vs Normal Gene Expression Profiles Identifies New Disease Entities and Common Transcriptional Programs in AML Patients.” Blood 123 (6): 894–904.

“Rectified Linear Units Improve Restricted Boltzmann Machines.” n.d. Accessed December 5, 2022. https://openreview.net/forum?id=rkb15iZdZB.

Schmauch, Benoît, Alberto Romagnoni, Elodie Pronier, Charlie Saillard, Pascale Maillé, Julien Calderaro, Aurélie Kamoun, et al. 2020. “A Deep Learning Model to Predict RNA-Seq Expression of Tumours from Whole Slide Images.” Nature Communications 11 (1): 3877.

Schuster, Viktoria, and Anders Krogh. 2021. “A Manifold Learning Perspective on Representation Learning: Learning Decoder and Representations without an Encoder.” Entropy 23 (11). https://doi.org/10.3390/e23111403.

Schuster, Viktoria, and Anders Krogh. 2022. “The Deep Generative Decoder: MAP Estimation of Representations Improves Modeling of Single-Cell RNA Data.” arXiv, November. https://doi.org/10.48550/arXiv.2110.06672.

Vaca-Paniagua, Felipe, Rosa María Alvarez-Gomez, Hector Aquiles Maldonado-Martínez, Carlos Pérez-Plasencia, Veronica Fragoso-Ontiveros, Federico Lasa-Gonsebatt, Luis Alonso Herrera, et al. 2015. “Revealing the Molecular Portrait of Triple Negative Breast Tumors in an Understudied Population through Omics Analysis of Formalin-Fixed and Paraffin-Embedded Tissues.” PLOS ONE. https://doi.org/10.1371/journal.pone.0126762.

Vihinen, Mauno. 2022. “Individual Genetic Heterogeneity.” https://doi.org/10.3390/genes13091626.

Vivian, John, Jordan M. Eizenga, Holly C. Beale, Olena M. Vaske, and Benedict Paten. 2020. “Bayesian Framework for Detecting Gene Expression Outliers in Individual Samples.” JCO Clinical Cancer Informatics 4 (February): 160–70.

Way, Gregory P., and Casey S. Greene. 2018. “Extracting a Biologically Relevant Latent Space from Cancer Transcriptomes with Variational Autoencoders.” Pacific Symposium on Biocomputing. Pacific Symposium on Biocomputing 23: 80–91.

Wilks, Christopher, Shijie C. Zheng, Feng Yong Chen, Rone Charles, Brad Solomon, Jonathan P. Ling, Eddie Luidy Imada, et al. 2021. “recount3: Summaries and Queries for Large-Scale RNA-Seq Expression and Splicing.” Genome Biology 22 (1): 323.

Zeng, William Z. D., Benjamin S. Glicksberg, Yangyan Li, and Bin Chen. 2019. “Selecting Precise Reference Normal Tissue Samples for Cancer Research Using a Deep Learning Approach.” BMC Medical Genomics 12 (Suppl 1): 21.

